# Phenotyping heart failure using model-based analysis and physiology-informed machine learning

**DOI:** 10.1101/2021.03.03.433748

**Authors:** Edith Jones, E. Benjamin Randall, Scott L. Hummel, David Cameron, Daniel A. Beard, Brian E. Carlson

## Abstract

To determine the underlying mechanistic differences between diagnoses of Heart Failure (HF) and specifically heart failure with reduced and preserved ejection fraction (HFrEF & HFpEF), a closed loop model of the cardiovascular system coupled with patient specific transthoracic echocardiography (TTE) and right heart catheterization (RHC) measures was used to identify key parameters representing cardiovascular hemodynamics. Thirty-one patient records (10 HFrEF, 21 HFpEF) were obtained from the Cardiovascular Health Improvement Project (CHIP) database at the University of Michigan. Model simulations were tuned to match RHC and TTE pressure, volume and cardiac output measures in each patient with average error between data and model of 4.87 ± 2%. The underlying physiological model parameters were then plotted against model-based norms and compared between the HFrEF and HFpEF group. Our results confirm that the main mechanistic parameter driving HFrEF is reduced left ventricular contractility, while for HFpEF a much wider underlying phenotype is presented. Conducting principal component analysis (PCA), *k*-means, and hierarchical clustering on the optimized model parameters, but not on clinical measures, shows a distinct group of HFpEF patients sharing characteristics with the HFrEF cohort, a second group that is distinct as HFpEF and a group that exhibits characteristics of both. Significant differences are observed (*p*-value<.001) in left ventricular active contractility and left ventricular relaxation, when comparing HFpEF patients to those grouped as similar to HFrEF. These results suggest that cardiovascular system modeling of standard clinical data is able to phenotype and group HFpEF as different subdiagnoses, possibly elucidating patient-specific treatment strategies.

## Introduction

Heart Failure with preserved ejection fraction (HFpEF) is diagnosed in patients with the hallmarks of heart failure and a left ventricular ejection fraction (EF) equal to or above 50%. HFpEF now represents more than half of heart failure cases and its incidence is increasing with an aging population and a high prevalence of associated risk factors (e.g., systemic hypertension, coronary artery disease, diabetes present in 75-90% of patients)(Yancy *et al*., 2006; Owan *et al*., 2006; Hummel *et al*., 2009; Little & Zile, 2012). Patients with HFpEF suffer low quality of life and poor long-term outcomes. Despite this substantial individual and public health burden, HFpEF lacks a framework for evidence-based pharmacotherapy(Yancy *et al*., 2013). Long-term management of HFpEF focuses on the treatment of any existing comorbidities, therapeutics that decrease the left ventricular diastolic pressures, and general symptom reduction. Several clinical trials in large cohorts of HFpEF patients have failed to demonstrate consistent benefit. The drugs used include sildenafil (Guazzi *et al*., 2011; Borlaug *et al*., 2015; Hoendermis *et al*., 2015; Liu *et al*., 2017), sacubitril/valsartan (Solomon *et al*., 2019), losartan (Wachtell *et al*., 2010), candesartan (Yusuf *et al*., 2003), spironolactone(Edelmann *et al*., 2013; Cohen *et al*., 2020), and isosorbide mononitrate(Redfield *et al*., 2015). Studies using these drugs have had some success but often trials using the same drug contradict each other (e.g. sildenafil studies).

HFpEF was previously termed “diastolic” heart failure, with symptoms attributed to ventricular stiffness and/or impaired relaxation, impaired ventricular filling during diastole, and higher average pressures during the cardiac cycle. However, patients with HFpEF have dysfunction in multiple cardiovascular domains, some of which may become evident only during exercise(Dunlay *et al*., 2017). It has been suggested that selecting the correct HFpEF cohort is an important factor in the study success(Borlaug *et al*., 2015), but the wide range of HFpEF phenotypes at the mechanistic cardiovascular systems level makes selecting these cohorts from upper level clinical measures difficult. Since HFpEF is a catch-all category for heart failure patients based mainly on EF estimates, the inability to have a standard treatment for these patients may be an indicator of the physiological heterogeneity underlying HFpEF. Therefore, identifying subpopulations of HFpEF patients with similar cardiovascular etiologies is a crucial task required to target appropriate therapies for these patients.

Patients presenting with heart failure and an EF below 50% are diagnosed with heart failure with reduced ejection fraction (HFrEF). The classical understanding of HFrEF, also known as “systolic” heart failure, is that ventricular dilation and reduced systolic contractility cause reduced ability to pump blood to the systemic circulation during systole (Pinilla-Vera *et al*., 2019). Unlike HFpEF, current therapies have been successfully applied to HFrEF patients, including ACE-inhibitors and Angiotensin II receptor blockers (ARBs) to modulate the renin-angiotensin system.

To diagnose and monitor patients with heart failure, two clinical procedures are commonly used: transthoracic echocardiography (TTE) and right heart catheterization (RHC). TTE is non-invasive, widely available, and may be used to quantify left ventricular volumes in systole and diastole for EF estimation. From TTE measurements, we may obtain additional information, such as cardiac output based on the heart rate and the left ventricular out track flow velocity time integral (LVOT VTI) for each patient. RHC is used to measure right ventricular and pulmonary artery pressures during systole and diastole along with CO, heart rate, and pulmonary capillary wedge pressure. While TTE and RHC provide detailed ventricular volume and pressure data for individual patients, the challenge of integrating these measures into a single representation of a patient’s cardiovascular state is only made qualitatively in the clinic. One way to quantitatively reconcile what these datasets show about the right and left sides of the heart is with a closed loop model of the cardiovascular system. To combine these two sets of data, we must take into consideration that the two datasets are typically not obtained simultaneously and may include single measures and time course data.

In this retrospective study, we have developed a methodology to represent the cardiovascular state of both HFpEF and HFrEF patients in an effort to determine the underlying mechanistic differences between diagnoses and specifically within the diagnosis of HFpEF. Recent studies have determined subpopulations of the HFpEF diagnosis using RNA sequencing (Hahn *et al*., 2021) and quantitative echocardiography (Shah, 2019). Here, we aim to discern subgroups within our HFpEF cohort using a modeling and unsupervised machine learning approach. To this end, a clustering analysis is performed on estimated model parameters identifying HFpEF subgroups that provide hemodynamic insight into functional differences between HFpEF patient subgroups. To our knowledge, this is the first attempt at categorizing HFpEF using physiologically based computational models of the cardiovascular system. A workflow of the approach used in this study is shown in **Figure 1**.

**Figure 1.**
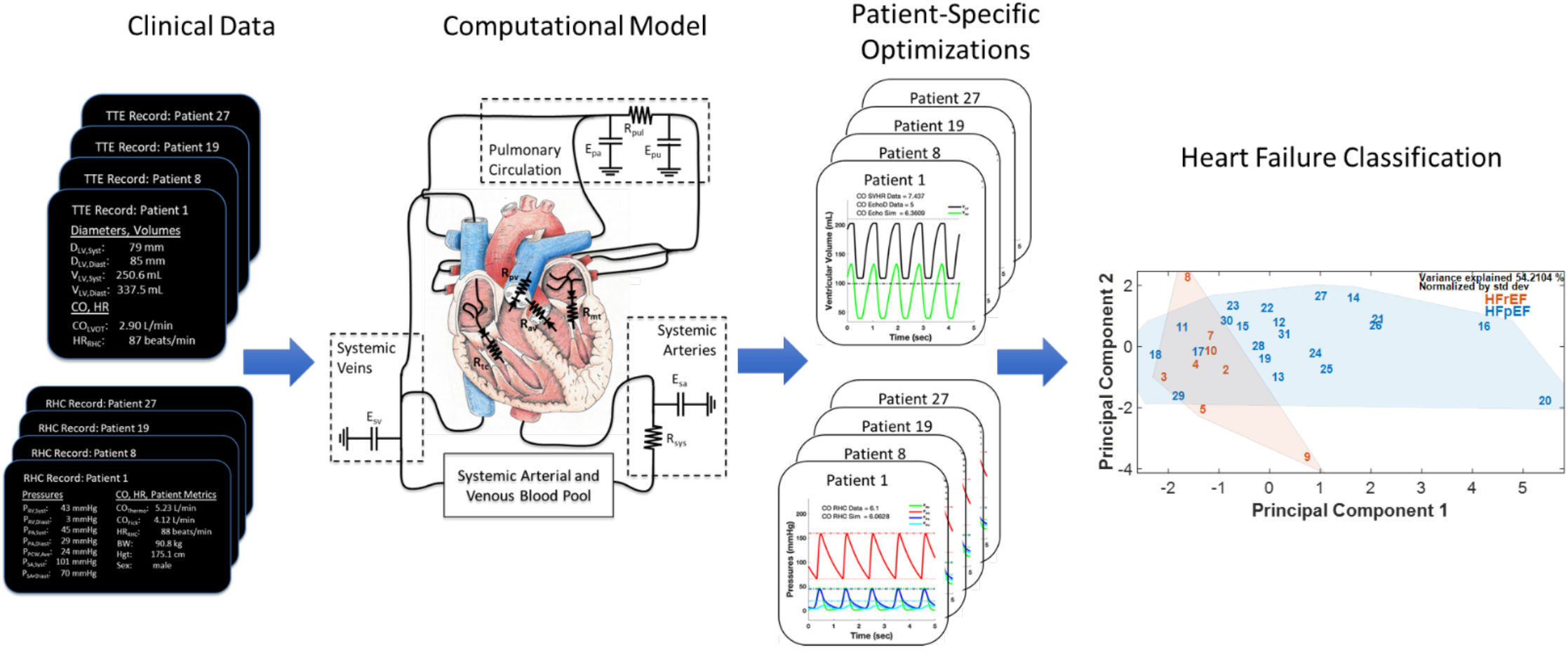
Approach for analysis of right heart catheterization (RHC) and transthoracic echocardiography (TTE) measures from patients with heart failure with preserved ejection fraction (HFpEF) and heart failure with reduced ejection fraction (HFrEF) using patient-specific cardiovascular system models. RHC and TTE measures are reproduced for each patient with the model by adjusting mechanistic parameters in the model. These underlying parameter values can then be used to observe differences between HFpEF and HFrEF patients and uncover subcategories of HFpEF using clustering methods.

## Methods

### Clinical Data

The Cardiovascular Health Improvement Project (CHIP) database was queried to extract clinical measures from patients diagnosed with HFpEF or HFrEF. This repository is a research-ready biorepository of DNA, plasma, serum, and tissue samples, which also includes de-identified electronic health records (EHRs) from consenting patients with heart failure, aortic disease, arrhythmia, and dyslipidemia. The CHIP repository is supported by the Frankel Cardiovascular Center at the University of Michigan. Through the CHIP office, a search was made to collect clinical measures from HFpEF and HFrEF patients with both a TTE and RHC contained within their EHR. The patients with both procedures were extracted from all HFpEF and HFrEF records in a time range from February 2016 through February 2019 and the time between RHC and TTE procedures was constrained to be less than 90 days. With this query, 62 patient records (26 HFrEF and 36 HFpEF) were collected. Patient records missing the minimal number of measures (see below) from RHC and TTE procedures eliminated 10 HFrEF and 13 HFpEF records, leaving 34 patient records (11 HFrEF and 23 HFpEF). Finally, one HFrEF and two HFpEF records that appeared to be outliers during the initial phase of our analysis were followed up in the patient record and were found to have procedures or treatments that changed their original cardiovascular diagnosis (e.g. chemotherapy changing a patient from HFpEF to HFrEF). These three patients were omitted from our final retrospective cardiovascular systems analysis leaving 31 patient records (10 HFrEF and 21 HFpEF).

#### RHC measures

During this invasive procedure, a Swan-Ganz catheter is inserted through the jugular vein and measures the pressure at the tip of the catheter as it is advanced into the pulmonary artery. Besides pressure information, CO is estimated by using the thermodilution or Fick methods. The thermodilution technique estimates CO by measuring dispersion of a cold saline bolus injected at the proximal end and then sensed at the distal end of the catheter. The Fick method measures venous and arterial O_2_ saturation and often assumes a given whole body oxygen consumption (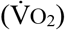) based on weight, height, and sex. The accuracy of the Fick method hinges on correctly estimating 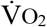, and it has been determined that it can vary by as much as 25% when compared to a direct measurement of 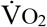 (Narang *et al*., 2014). Both direct 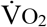 and estimated 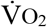 are used in the RHC lab at the University of Michigan, however all RCH records in this study used estimated 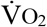, therefore we have chosen the thermodilution method as a consistent measure of RHC CO.

The selected RHC datasets came from reports that contained at least the following thirteen clinically measured values: systolic and diastolic right ventricular pressure, systolic and diastolic pulmonary arterial pressure, average pulmonary capillary wedge (PCW) pressure, systolic and diastolic systemic pressure, heart rate during the RHC, CO (thermodilution and Fick), body weight, height, and sex as shown in **Table 1**. In an attempt to ensure that the measures are taken as close together as possible, heart rate and systolic and diastolic systemic pressure were gathered from the RHC report only while the catheter was inserted. If multiple measures were taken during this period, an average was taken of the values recorded.

**Table 1:**
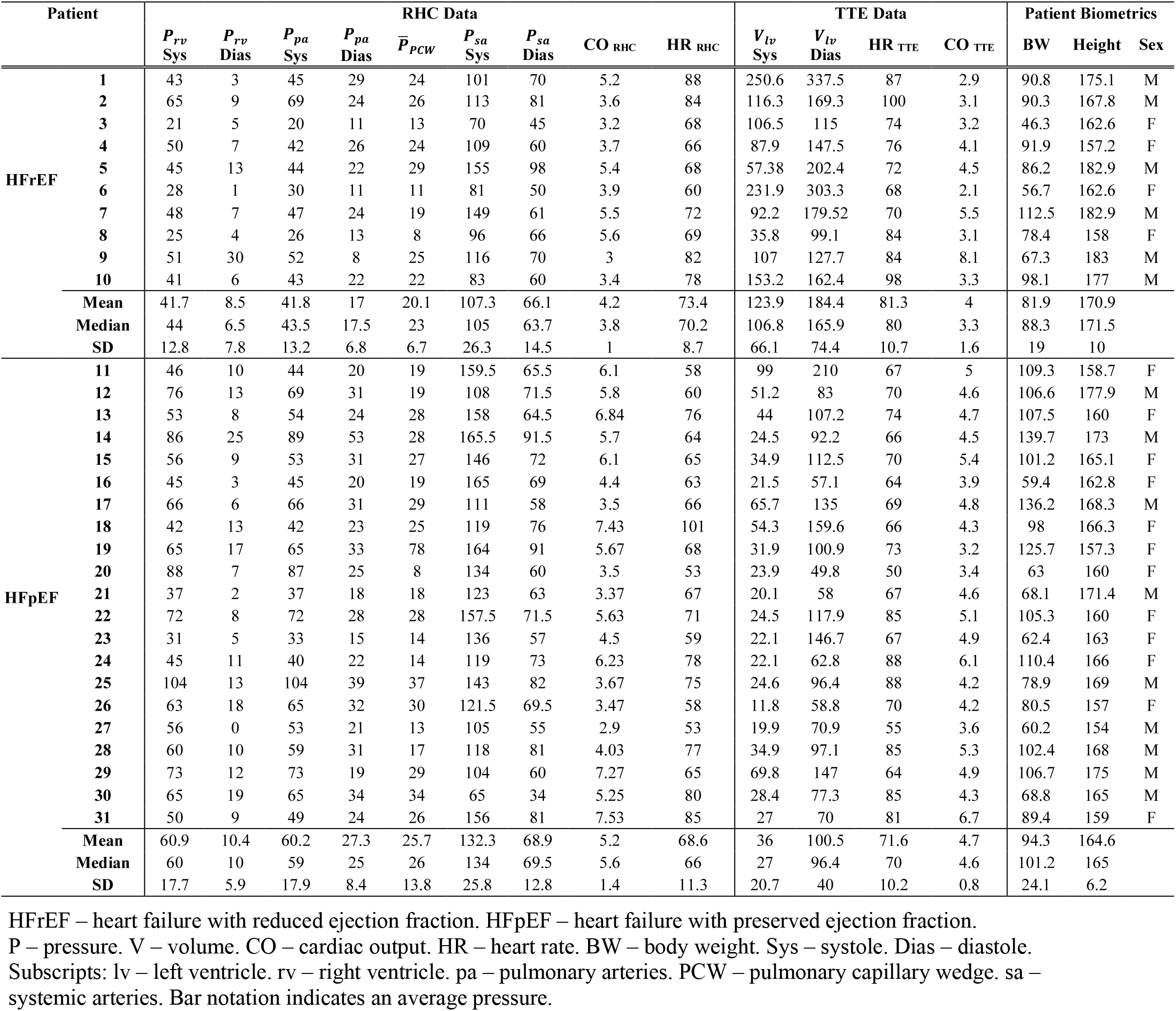
Right heart catheterization (RHC) data, transthoracic echocardiogram (TTE) data, and patient biometrics.

#### TTE measures

The selected TTE datasets include at a minimum: measurements of left ventricular volume in systole and diastole and heart rate during TTE. The left ventricular volumes in systole and diastole are measured as either a single diameter across the left ventricle just below the mitral valve leaflet tips or volumes quantified from tracings of the left ventricle from apical two- and four-chamber views(Lang RM *et al*., 2016). The single diameter derived volumes assume the left ventricle can be approximated as a truncated prolate spheroid with a nonlinear relationship between the diameter and length of the ventricle (Narang *et al*., 2014). Volumes derived from the two- and four-chamber views are calculated by the method of discs, also known as Simpson’s method (Opotowsky *et al*., 2017). Since Simpson’s method is preferred for the estimation of left ventricular volumes over the single diameter estimation, all patient raw TTE images were reviewed by a cardiologist to (i) obtain a Simpson’s method estimate when the quality of the image allowed, (ii) determine the heart rate, and (iii) extract an LVOT VTI estimate of CO when possible. This resulted in a possibility of having three separate estimates of CO: heart rate times the stroke volume by single diameter volume estimates, heart rate times the stroke volume by Simpson’s method, and heart rate times LVOT VTI times the cross sectional area of the outflow tract(Lang RM *et al*., 2016). Thus, it is necessary to create the systematic method of ranking the quality of these measures and determining a CO to be used for parameter optimization in the patient-specific cardiovascular systems model described below.

#### Decision trees

Both TTE and RHC measures can contain multiple estimates of CO. Similarly, TTE itself can contain multiple estimates of left ventricular volume in systole and diastole. Therefore, we have developed a consistent method to determine whether to select one of the estimates or average them, as shown by the decision trees (**Figure 2**).

**Figure 2.**
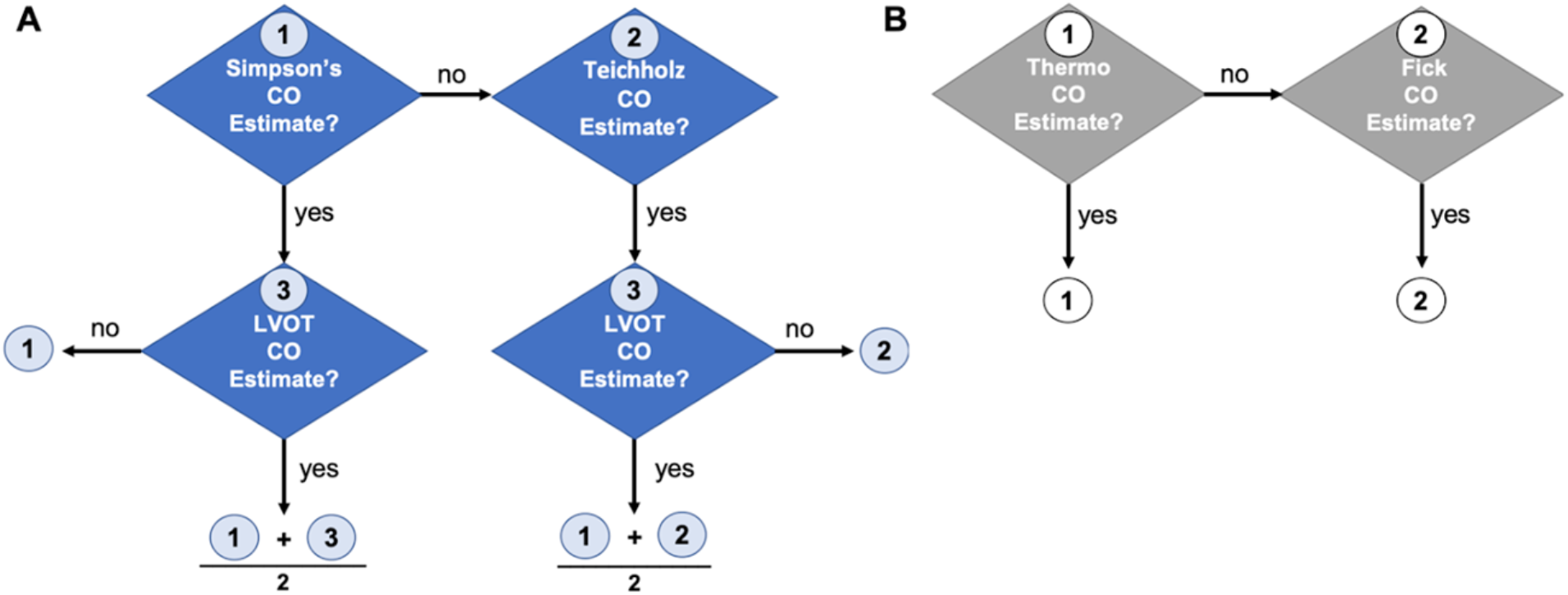
Decision trees to determine which cardiac output estimates to use in patient data records for TTE and RHC. **A**. For TTE, cardiac output estimates from LV volumes in systole and diastole through the Simpson’s method (1) was the first choice used in our study. If Simpson’s method was not available cardiac output estimates would be conducted from LV diameter data during systole and diastole using Teichholz formula (2). If an LVOT VTI cardiac output estimate (3) was also available, it was averaged with either the Simpson’s or single diameter cardiac output estimate **B**. cardiac output estimate used in RHC gives priority to thermodilution estimates as explained previously.

The decision tree for the TTE has two cases (**Figure 2A**). Case 1: if the Simpson’s method estimate is available, it takes priority and is used for CO and left ventricular volume estimates. Case 2: if the two- and four-chamber images do not (Teichholz *et al*., 1976) yield a Simpson’s method estimate, the volumes and CO are calculated from single diameter measures using the Teichholz equation. In both cases, if there is a LVOT VTI estimate, the LVOT VTI estimate is averaged with the CO estimate. If not, the initial estimate is taken as the CO for the patient.

In estimating CO from the RHC measurements, if the thermodilution CO estimate is available, it is used. If not, CO is calculated via the Fick method, as shown in **Figure 2B. Table 1** lists the data used in this study screened with these decision criteria.

### Mathematical modeling framework

The cardiovascular systems model is similar to that used in a previous study from our lab (Colunga *et al*., 2020) and is based on the formulation developed by Smith et al. (Smith *et al*., 2004). The model complexity was reduced significantly since the clinical data used for parameterization here do not have enough informational content to uniquely identify the parameters of the full Smith et al. model. In our previous reduced version of the model, ventricular-ventricular interaction and fluid inertance after each heart valve were omitted from the Smith et al. model. In addition, for this study, the pericardial compartment was removed and the zero pressure (or dead space) volumes in all vascular and ventricular compartment were set to zero. We have added a factor to recruit stressed blood volume from the nominal value of 30% total blood volume, which has been posited to occur in heart failure (Fudim *et al*., 2017); however, no volume was recruited to represent any of the HFpEF or HFrEF patients in this study. Equations for the reduced cardiovascular system model used in this study are given in Appendix A and model code without parameter optimization can be found on GitHub.

#### Nominal parameters and initial conditions

The reduced cardiovascular system model used in this study has 16 model parameters that can potentially be adjusted during the optimization procedure. Parameter descriptions are listed in **Table 2**. Nominal estimates of all parameters are determined starting with the set of expressions from our previous study (Colunga *et al*., 2020) as a guide. However, some assumptions used in the previous nominal parameter calculations can be replaced with data since TTE measurements are available. Therefore, a reformulation of some nominal parameter expressions has been made in this study:

**Table 2:** Model parameters for patients at rest.

| Parameter |  | Units | Description | Fixed | Adjustable |
| --- | --- | --- | --- | --- | --- |
| **Left Ventricle (LV)** | $E_{lv}$ | mmHg mL<sup>-1</sup> | LV active contractility | | X |
| | $P_{0,lv}$ | mmHg | LV diastolic reference pressure | X | |
| | $\lambda_{lv}$ | mL<sup>-1</sup> | LV passive stiffness | | X |
| **Right Ventricle (RV)** | $E_{rv}$ | mmHg mL<sup>-1</sup> | RV active contractility | | X |
| | $P_{0,rv}$ | mmHg | RV diastolic reference pressure | X | |
| | $\lambda_{rv}$ | mL<sup>-1</sup> | RV passive stiffness | | X |
| **Pulmonary Vasculature** | $E_{pa}$ | mmHg mL<sup>-1</sup> | Pulmonary arterial stiffness | | X |
| | $E_{pv}$ | mmHg mL<sup>-1</sup> | Pulmonary venous stiffness | | X |
| | $R_{pul}$ | mmHg s mL<sup>-1</sup> | Pulmonary resistance | | X |
| **Systemic Vasculature** | $E_{sa}$ | mmHg mL<sup>-1</sup> | Systemic arterial stiffness | | X |
| | $E_{sv}$ | mmHg mL<sup>-1</sup> | Systemic venous stiffness | X | |
| | $R_{sys}$ | mmHg s mL<sup>-1</sup> | Systemic resistance | | X |
| **Heart Valves** | $R_{mt}$ | mmHg s mL<sup>-1</sup> | Mitral valve resistance | X | |
| | $R_{av}$ | mmHg s mL<sup>-1</sup> | Aortic valve resistance | X | |
| | $R_{tc}$ | mmHg s mL<sup>-1</sup> | Tricuspid valve resistance | X | |
| | $R_{pv}$ | mmHg s mL<sup>-1</sup> | Pulmonary valve resistance | X | |

**Table 3:**
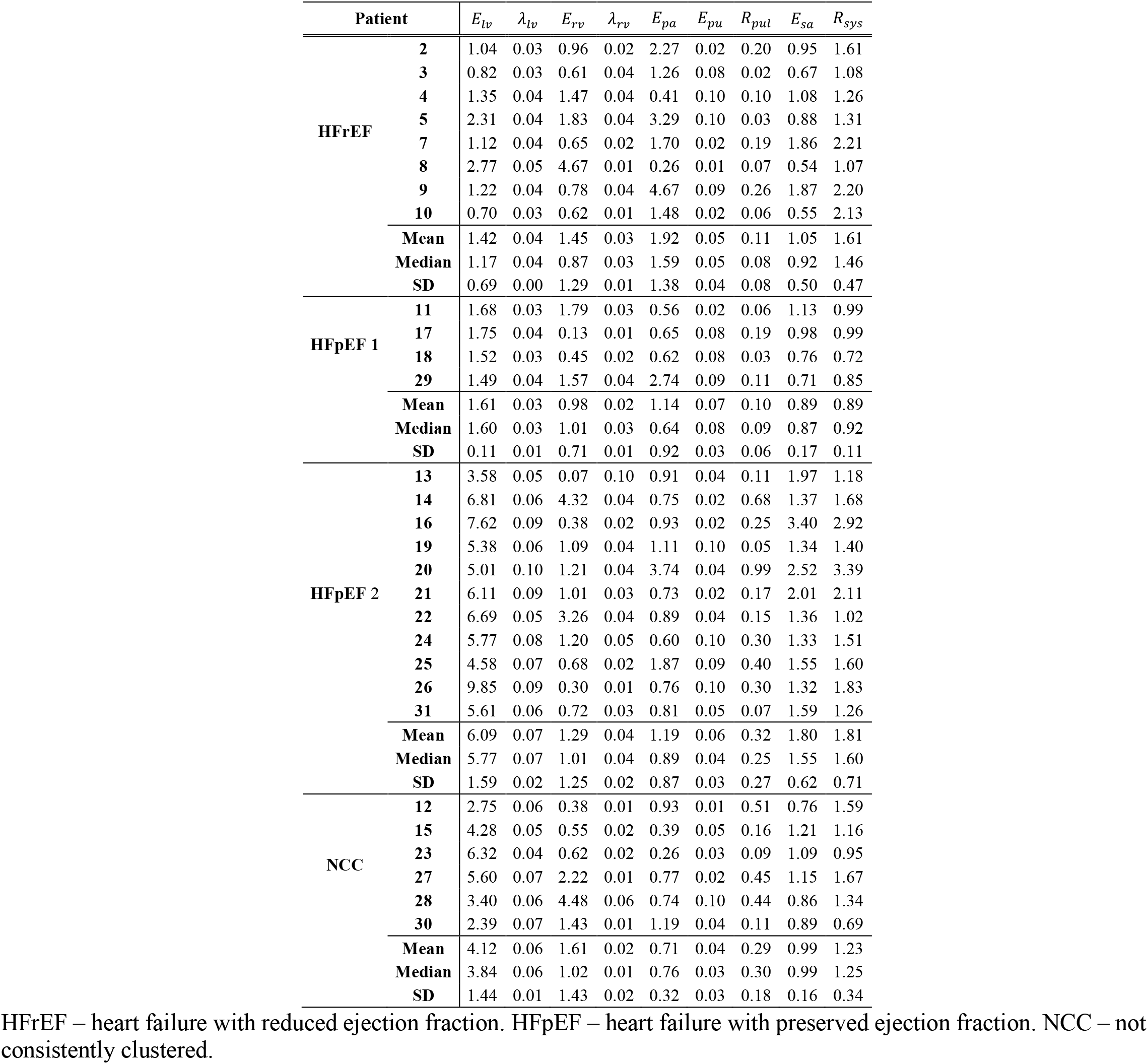
Patient specific optimized parameter values.

- left ventricular elastances were calculated from measured volume and estimated pressure in systole;
- left ventricular diastolic stiffness was calculated from measured volume and estimated pressure in diastole;
- right ventricular elastances were calculated from measured pressure and estimated volume in systole;
- right ventricular diastolic stiffness was calculated from measured pressure and estimated volume in diastole;
- systemic elastances were calculated from measured pulse pressures for the arterial compartments, estimated pulse pressures for the venous compartments, and estimated stressed volumes;
- pulmonary elastances were calculated from estimated and measured pulse pressures and estimated stressed volumes;
- systemic and pulmonary resistances were calculated from measured systemic and pulmonary average arterial pressures and estimated systolic venous pressure along with the measured RHC cardiac output.

Resistances across the four valves were calculated in exactly the same way as in our previous study and the ventricular end diastolic reference pressures were set to normal values from Smith et al. More details on the exact expressions used for nominal calculations are shown in Supplemental Material and the model code on GitHub.

Total blood volume is calculated based on the height, weight, and sex of each patient as described in our study looking at heart transplant patients (Colunga *et al*., 2020), utilizing the expression originally developed by Nadler et al. (Nadler *et al*., 1962) The initial distribution of stressed and unstressed blood volume among the six vascular compartments is based on the work by Beneken (Beneken, 1968), in which a total stressed volume of 18.75% was assumed. In this study, we assumed 30% of the total blood volume is stressed volume (Beneken, 1968; Fudim *et al*., 2017; Colunga *et al*., 2020), so additional volume was recruited from the four systemic and pulmonary compartments based on the unstressed volume available in each compartment. Complete details of adapting the Beneken volume distributions for this study can be seen in Supplemental Material and the model code on GitHub.

#### Global sensitivity analysis

To determine which of these parameters can be identified with the given clinical patient data, a sensitivity analysis is performed. Due to the vast variation in parameter values across subjects, we conducted a global sensitivity analysis using Sobol’ indices to explore the entire parameter space. Sobol’ indices apportion the variance in the output to the effect of each parameter (Sobol′, 2001). In particular, we use total effect Sobol’ indices to characterize the effect of both the parameter and parameter interactions on the residual variance (Randall E. B. *et al*., 2021). This residual was calculated by determining the least square error between simulations and RHC and TTE data as described in our previous study (Colunga *et al*., 2020). We then ranked the total effect Sobol’ indices to determine a set of influential parameters that substantially affect the variance of the residual, i.e., a subset of parameters that have an index above the threshold *η* = 10^−3^. Parameters below the threshold were excluded from consideration for optimization and set to their nominal values. Though the parameters for diastolic pressure development in the right and left ventricle (*P*_*0,lv*_ and *P*_*0,rv*_) and elastance of the systemic veins (*E*_*sv*_) were above *η*, they cannot be determined explicitly (Colunga *et al*., 2020). Hence, *P*_*0,lv*_ and *P*_*0,rv*_ were set to the values used in Colunga et al., and *E*_*sv*_ was calculated nominally as described above. Note that our previous study used only RHC data to determine model parameters. Since TTE data were included here, two additional model parameters could be identified: the right and left ventricular free wall elastance, *E*_lv_ and *E*_rv_, respectively. From the set of influential parameters, we obtained the estimable subset

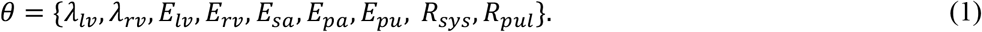

This adjustable parameter subset aligns well with the parameter subset used in our earlier work (Colunga *et al*., 2020).

#### Optimization

For each patient, we estimate the adjustable parameters in Equation (1) by minimizing the least square error between the simulations and data for ten measures: right ventricular pressure in systole and diastole, pulmonary artery pressure in systole and diastole, average pulmonary capillary wedge pressure, systemic artery pressure in systole and diastole, CO during RHC, left ventricular volume in systole and diastole, and CO during TTE. Since the heart rate during RHC and TTE can be different, two separate simulations are run – one simulating the RHC and one simulating the TTE; however, both simulations are run with one set of parameter values with the assumption that the parameters representing cardiac function do not change appreciably across procedures for a single patient. Values of the clinical measures are calculated over the cardiac cycle after the system has reached a steady state of pulsatile pressures and flows. This is assured by allowing our simulations to run for 50 beats. Once this steady state is reached, the maximum and minimum values of the pressure and volume data of the last 5 beats are used to compute the total residual error. The capillary wedge pressure and the CO represent average values over the cardiac cycle; because of this, their values are averaged over the cardiac cycle before being compared to the TTE and RHC measures. Estimates for the adjustable parameters are obtained using a genetic algorithm optimization implemented in Matlab (MathWorks Natick, Ma).

### Machine learning

We utilized three different clustering techniques to group individuals within a population based on similar characteristics. In theory, patients within the same groups should share similar physiological characteristics. The clinical data and optimized parameter values were compiled into separate matrices, *D* and *P*, respectively, where each row represents a given patient and each column represents a clinical measure or optimized parameter value (**Tables 1** and **2**). Before any of the clustering methods are applied, each column is centered by subtracting the average of each column from each element in that column. Because our clinical data and optimized parameters had different units within their respective matrices, we standardized each variable to unit norm by dividing each variable by its standard deviation.

#### Principal Component Analysis (PCA)

We performed a PCA (Jolliffe IT, 1986), which is simply a singular value decomposition identifying an orthogonal change of basis within the clinical data or optimized parameter spaces that retains the greatest variation across patients independent of level of dimension reduction selected. For the optimized parameter matrix *P*, the decomposition *P* = *USV*^′^ produces unitary matrices *U* and *V*and diagonal matrix *S*, representing the portion of the total variation explained by each principal component. The PCA score, which gives the position in this rotated space that maximizes variation, is given by the product of *U* and *S*. A convex hull was prescribed around both the HFpEF and HFrEF patients to delineate the regions occupied by the HFpEF and HFrEF clinical diagnosis groups. An example of the location of each patient and the prescribed regions considering only the first two principal components with respect to the clinical data is shown in **Figure 4A**. Patients that fall into the overlap region are considered ambiguous and assigned a group based on the following clustering methods.

#### k-means

*k* -Means clustering creates *k* unsupervised clusters from the data. In this study we chose to group the patients into two clusters, so two patients are randomly chosen as a cluster centroids, and all other patients are grouped based on which centroid patient they are closest to. The centroid of these new clusters is recalculated and the process is repeated until the cluster groupings do not change. We use the same centered and normalized vectors for each patient as with the PCA. This method is dependent on the random initial cluster centroids selected, so we run this process twenty times and select the clustering result that has the smallest total cluster variance (Eisen *et al*., 1998; Wilkin & Huang, 2008). **Figure 4B** shows two *k*-means clusters superimposed on the clinical measures PCA hulls.

#### Hierarchical clustering

Each patient starts as a cluster, and then hierarchical clustering finds the two closest patients and clusters them together. This process is repeated, grouping the two closest clusters together to reduce the total number of clusters by 1 until we have all the patients in one cluster(Kraskov *et al*., 2005). This method forms a hierarchical cluster tree known as a dendrogram that can then be truncated to produce the desired number of clusters. In Matlab, the linkage function uses the Ward metric (Ward, 1963), which groups the two clusters together that minimize the total in-cluster variation. Using the dendrogram, we partitioned our patients into two clusters by cutting the tree halfway between the second-from-last and last linkages. **Figure 4C** superimposes the hierarchical clusters on the PCA hulls.

Our focus is to identify groups that cluster consistently among these methods, especially since they use different concepts to group the data. If two patients share a PCA diagnosis, a *k*-means cluster, and a hierarchical cluster, they are grouped together. The clusters with the most HFrEF patients are considered the most “HFrEF-like”. Patients that switch between clusters for different methods are deemed not consistently clustered (NCC).

To classify the patients that fall in the PCA overlap region, we rely on the clustering methods. If a HFpEF patient in the overlap region falls in the *k*-means and hierarchical clusters that contain a majority of the HFrEF patients, we classify them as HFrEF-like HFpEF and thus are part of the HFpEF 1 group. Conversely, if they fall in the *k*-means and hierarchical clusters that contain a majority of HFpEF patients, we classify them as “pure” HFpEF and are part of the HFpEF 2 group. If they switch between clusters, they are classified as NCC.

## Results

Our retrospective cardiovascular systems analysis consists of a cohort of 31 patient records (10 HFrEF and 21 HFpEF). First, we consider the clinical data explicitly from the RHC and TTE **(Figure 3)**. Statistically significant differences in the means of TTE derived measurements (p-value < 0.001) such as ejection fraction, systolic & diastolic left ventricular volumes as well as cardiac output (p-value < 0.01) were found between HFrEF and HFpEF patients **(Figure 3A-D)**. Consistent with their systolic dysfunction phenotype, HFrEF patients had greater ventricular volumes than the HFpEF cohort, with patients 1 and 6 showing extreme ventricular dilation compared to the rest of the HFrEF cohort **(Figure 3 B, C)**. In the HFpEF cohort, pulmonary arterial systolic and diastolic pressures as well as right ventricular systolic pressures were significantly higher (p-value < 0.01) than the HFrEF cohort **(Figure 3 E-F)**. Systolic arterial pressure is likewise significantly higher (p-value < 0.05) in the HFpEF cohort when compared to HFrEFs (**Figure 3H)**.

**Figure 3.**
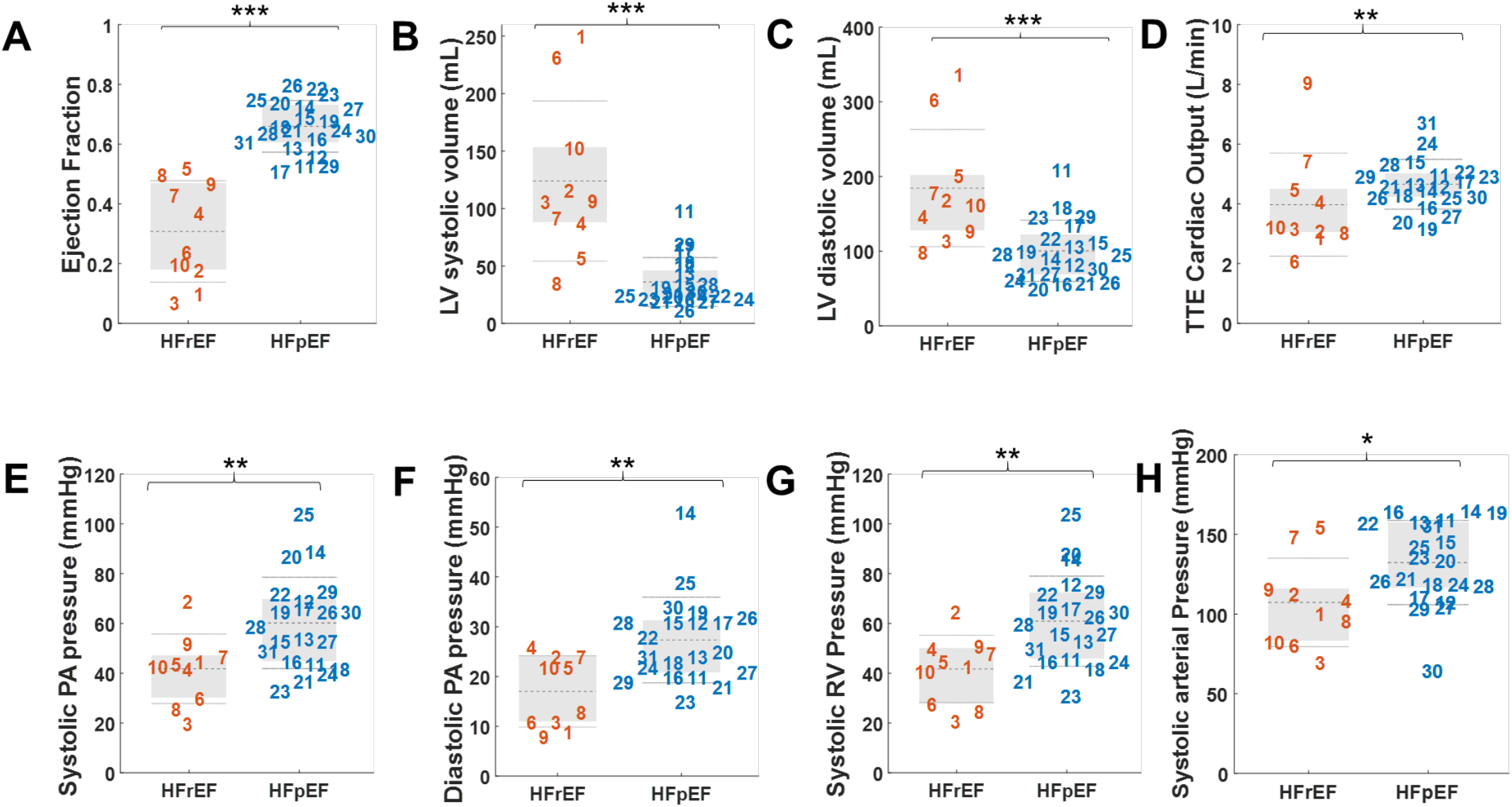
Significant differences are seen in TTE and RHC clinical data between heart failure patients based on their Ejection Fraction classification. Gray dashed line is average, light dotted is standard deviation and the grey box contains the middle 50% of the clinical values (* p-value <.05, ** p-value <.01, *** p-value <.001).

To determine if novel subpopulations of HF patients with similar cardiovascular etiologies could be discerned from clinical data alone, we performed a PCA along with two unsupervised clustering methods on the clinical data available from the RHC and TTE (**Figure 4)**. In **Figure 4A**, PCA scores for the first and second principal components are plotted and convex hulls are drawn around each group as they were clinically diagnosed. The convex hulls for HFrEF (orange) and HFpEF (blue) overlap, consisting of HFrEF patients 2,4, 5, 7, and 9 and HFpEF patients 11, 24, and 18. As PCA only captures the greatest spread across all patients, we are unable to determine whether the HFrEF and HFpEF patients in the overlap region share characteristics with the HFrEF or HFpEF clinical populations.

**Figure 4.**
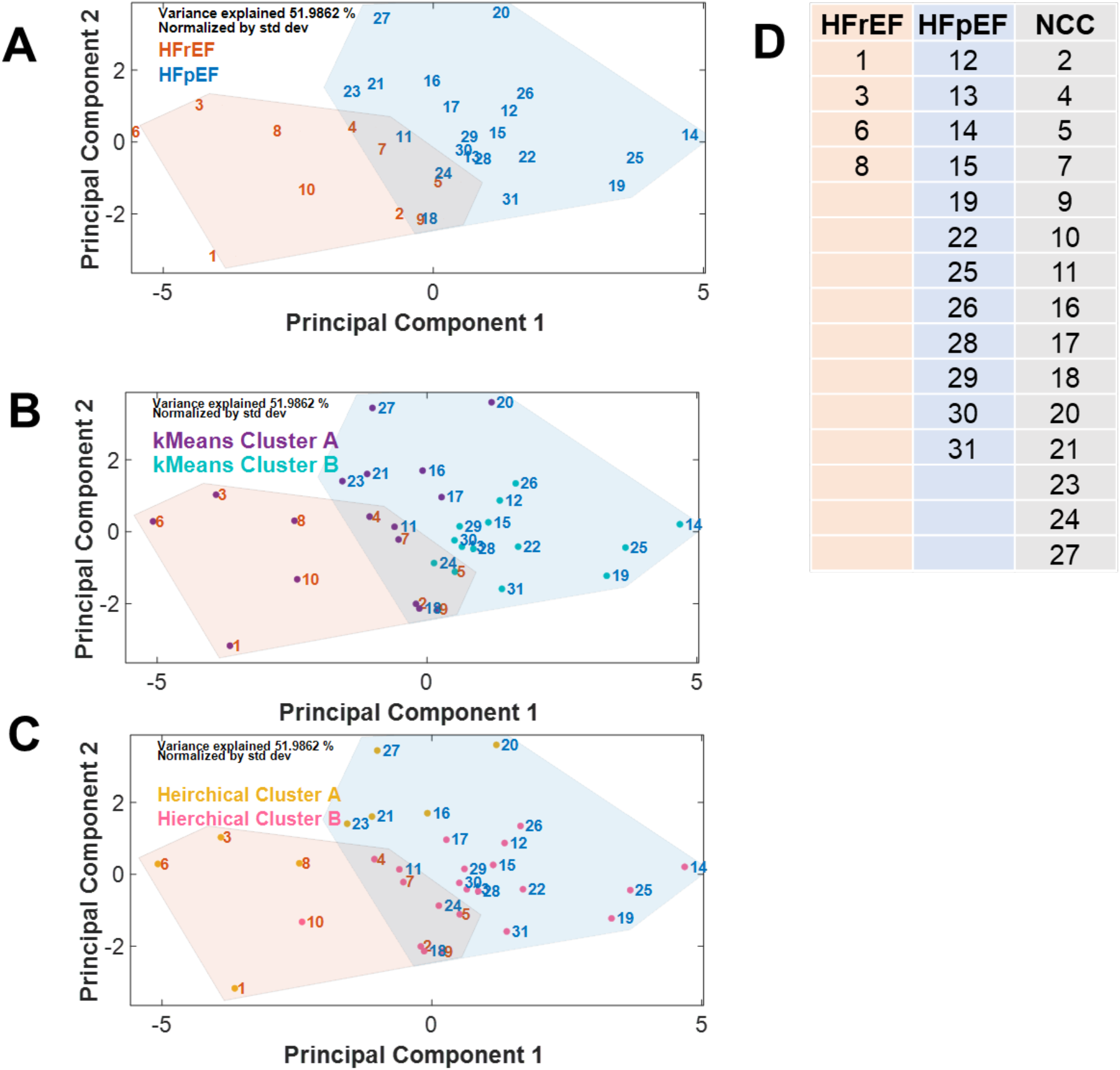
Clinical data clustering analysis **A**. Principal component analysis (PCA) of transthoracic echocardiogram (TTE) and right heart catheterization (RHC) clinical measurements. **B**. k-Means clustering of patient data with superimposed PCA where cluster A is more HFrEF-like and cluster B is more HFpEF-like. **C**. Hierarchical clustering of patient data with superimposed PCA analysis where cluster A is more HFrEF-like and cluster B is more HFpEF-like. **D**. Patient classification based on clinical measures clustering results. The HFrEF classification is for patients that fall in the HFrEF PCA hull, k-means cluster A, and hierarchical cluster A. The HFpEF classification is for patients that fall in the HFpEF PCA hull, k-means cluster B, and hierarchical cluster B. NCC refers to patients that are not consistently clustered among these methods.

To test if we could attribute the patients that fall in this PCA overlap region into a distinct phenotype associated with the HFrEF or HFpEF clinical groups, we employed k-means and hierarchical clustering (**Figure 4B, C**). We overlaid our k-means clustering results with our PCA convex hull (**Figure 4B)**. Since all HFrEF patients except patient 5 fall into k-means cluster A, we designate cluster A as more HFrEF-like and, conversely, k-means cluster B as more HFpEF-like. We observed in the overlap region, HFpEF patients 11 and 18 are in k-means cluster A whereas patient 24 is in k-means cluster B. Also, HFrEF patient 5 falls in k-means cluster B. Lastly, HFpEF patients 16, 17, 20, 21, 23, and 27 fall into k-means cluster A. In a similar fashion, hierarchical clustering results were overlaid with our PCA convex hull (**Figure 4C)**. Similarly, we specified hierarchical cluster A as more HFrEF-like and hierarchical cluster B as more HFpEF-like. Of particular interest is that all HFpEF patients in the overlap region now fall in hierarchical cluster B.

Among the clustering methods used here, **Figure 4D** denotes which patients consistently cluster in the following groups:

- “pure” HFrEF (n = 4) - patients that fall in the HFrEF PCA hull, k-means cluster A, and hierarchical cluster A;
- “pure” HFpEF (n = 12) - patients that fall in the HFpEF PCA hull, k-means cluster B, and hierarchical cluster B; and
- NCC (n = 15) - patients that do not consistently cluster.

**Supplemental Table 1** shows the patient designation based on clinical measurement clustering analysis. Note that through the use of a PCA, k-means and hierarchical clustering to analyze the clinical data available from the RHC and TTE, no subpopulations of HFpEF patients are revealed. These results also show many patients that fall in the NCC designation, both from the HFrEF and HFpEF diagnoses.

HFrEF subjects 1 and 6 show extreme ventricular dilation compared to other HFrEF patients in our cohort, with their systolic and diastolic volumes falling out as outliers in our whisker-plots (**Figure 3B, C)**. Because clinical data analysis indicated HFrEF 1 and 6 to be HFrEF outliers, these patients were excluded from further analysis. Moreover, when subjecting the clinical measures of these patients to clustering analysis, it was observed regardless of the specific clustering technique employed HFrEF patients 1 and 6 are found to be outside of the HFrEF hull (**Figure 4A)**.

To learn about the underlying physiological differences between our patient cohorts that cannot be determined from clinical data alone, patient clinical measurements were used to parameterize a reduced version of the Smith et al model. We conducted a global sensitivity analysis exploring the entire permissible parameter space and ranked the parameters due to their contribution to the residual. **Figure 5** displays the ranked total Sobol’ indices for all 16 adjustable parameters. This analysis shows that 12 parameters are influential to the residual, and from these parameters, we selected a subset of parameters to optimize, given in **Equation (1)**.

**Figure 5.**
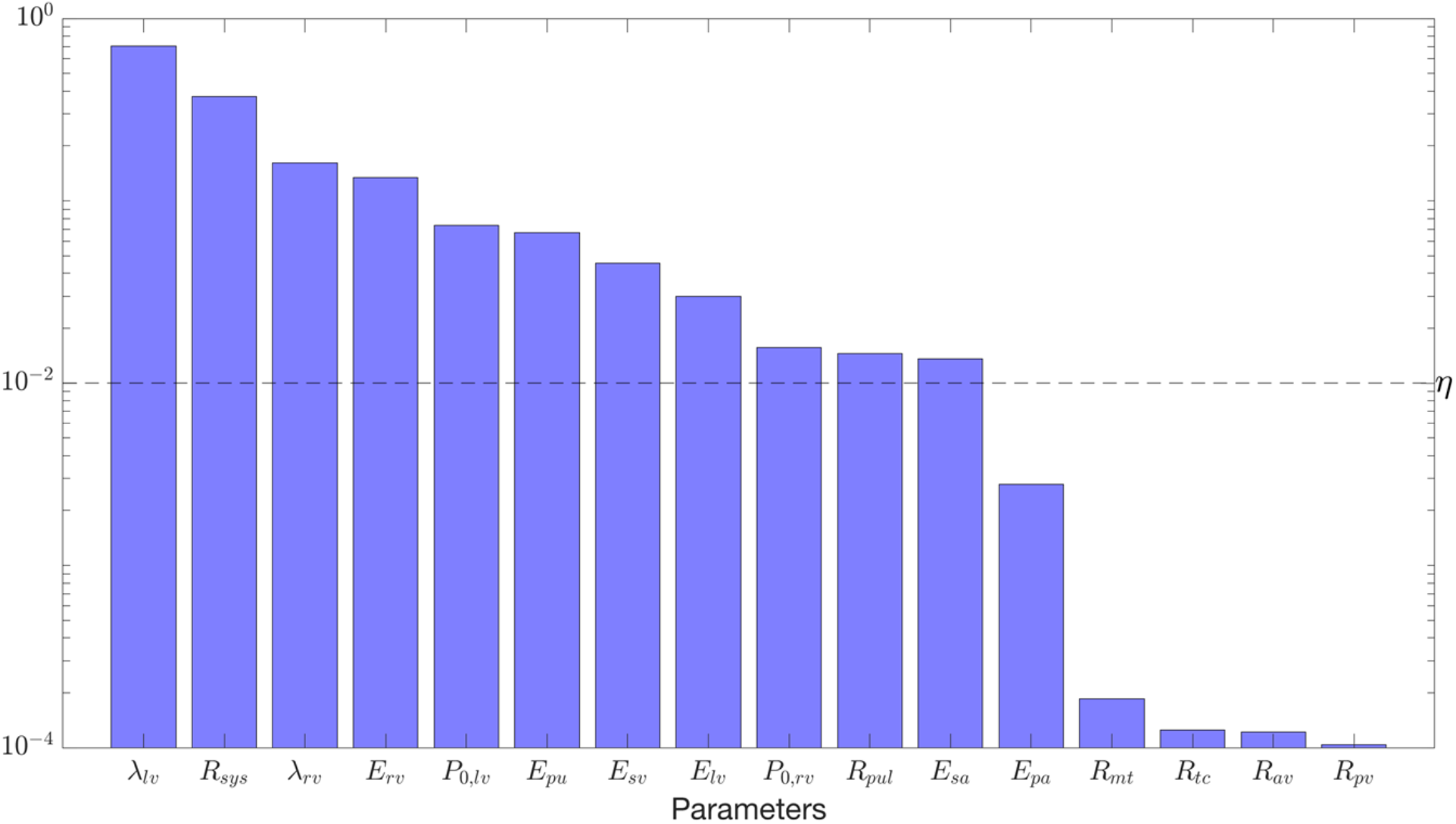
Global Sensitivity Analysis. Ranked total Sobol’ indices for all 16 adjustable parameters with an index above the threshold *η=10-3* were plotted with a vertical y-axis. This analysis shows that 12 parameters are influential to the residual. From these parameters, we selected a subset of parameters to optimize, given in **Equation (1)**.

Our model simulations predict that the HFpEF cohort has a much wider distribution of phenotype drivers than the cohort of HFrEF patients. Thus, we performed the same methods applied to the clinical measures to the optimized parameter values to identify subpopulations of HFpEF patients with similar cardiovascular etiologies **(Figure 6)**. The PCA scores of optimized parameters reveal a subpopulation of the heterogenous HFpEF cohort clustered by the HFrEF group, i.e., HFpEF patients 11, 17, 18 and 29 fall in the PCA overlap region **(Figure 6A)**.

**Figure 6.**
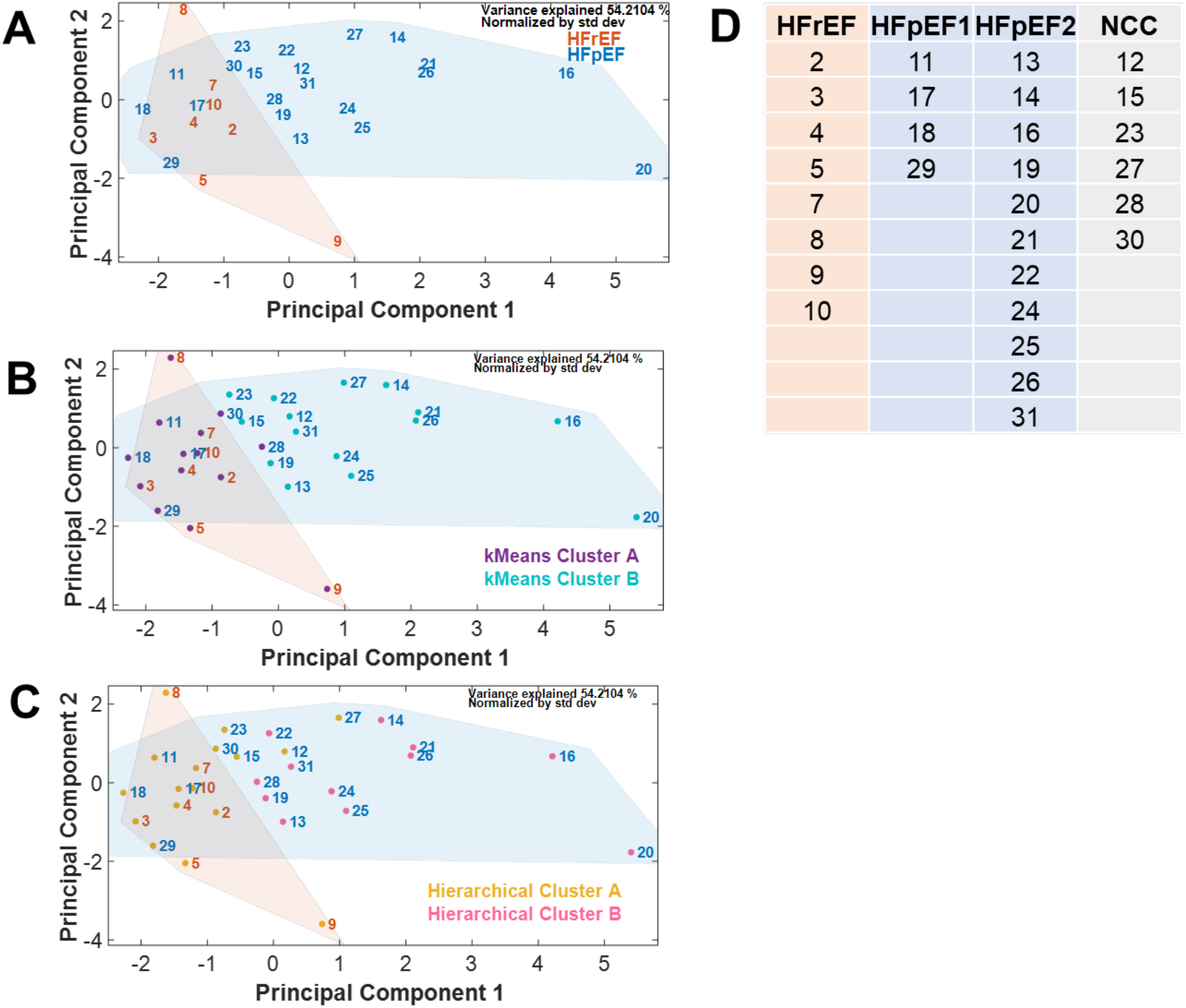
Clustering analysis of optimized model parameters shows three distinct groups of HFpEF patients. **A**. PCA analysis of optimized model parameters **B**. k-Means clustering of patients based optimized model parameters with superimposed PCA analysis. **C**. Hierarchical clustering of patients based on optimized model parameters with superimposed PCA analysis. **D**. Patient classification based on optimized model parameters clustering results. The HFrEF classification is for patients that fall in the HFrEF PCA hull, k-means cluster A, and hierarchical cluster A. The HFpEF Group 1 classification is for patients that fall in the HFpEF PCA hull, k-means cluster A, and hierarchical cluster A. The HFpEF Group 2 classification is for patients that fall in the HFpEF PCA hull, k-means cluster B, and hierarchical cluster B. NCC refers to patients that are not consistently clustered among these methods.

To determine whether this HFpEF subpopulation sharing strong patterns with the HFrEF cohort is distinct from the other HFpEF patients, we conducted k-means (**Figure 6B**) and hierarchical (**Figure 6C**) clustering on the optimized parameters that reveals a much different structure than clustering based on raw clinical data alone. In both clustering methods, all of the HFrEF patients fell into one cluster, which we designate as cluster A. Notably, all ambiguous patients also fall into cluster A for both methods; therefore, we conclude that this is a distinct HFpEF subpopulation.

The chart in **Figure 6D** indicates that conducting an independent analysis with PCA, k-means, and hierarchical clustering on the optimized model parameters reveals that out of the 29 patients selected for this final retrospective cardiovascular systems study fall into distinct groups:

- “pure” HFrEF (n=8) - patients that fall in the HFrEF PCA hull, k-means cluster A, and hierarchical cluster A;
- HFpEF Group 1 (n = 4) - patients that fall in the PCA overlap region, k-means cluster A, and hierarchical cluster A;
- HFpEF Group 2 (n = 11) - patients that fall in the HFpEF PCA hull, k-means cluster B, and hierarchical cluster B; and
- NCC (n = 6) - patients that do not consistently cluster.

**Supplemental Table 2** shows the patient designation based on optimized parameter clustering analysis. Since HFpEF Group 1 shares most all of the characteristics of the pure HFrEF group, we consider this group as more HFrEF-like, whereas HFpEF Group 2 is “purely” HFpEF. Notably all of the patients belonging to the not consistently clustered NCC group are HFpEF patients.

**Figure 7** illustrates the patient-specific values of key model parameters representing left ventricular (LV) active contractility, LV passive stiffness, systemic arterial stiffness, systemic and pulmonary resistance. Specifically, when comparing LV active contractility **(Figure 7A)** very significant differences are found amongst the different groups. Yet some trends are followed in some groups, with HFrEF and HFpEF Group 1 found to be below threshold and HFpEF Group 2 and NCC above threshold when compared to model-based norms **(Table 4)**. When compared to both HFrEF and HFpEF Group 1, the HFpEF Group 2 shows significantly higher LV active contractility (p-value < 0.001). Likewise, the NCC LV active contractility is significantly higher (p-value < 0.001 and p-value < 0.05, respectively) when compared to HFrEF and HFpEF Group 1. Of note no significant differences were found between HFrEF and HFpEF Group 1 in this parameter, and despite having similar trends HFpEF Group 2 had a significantly higher (p-value < 0.05) LV active contractility when compared to the NCC subgroup.

**Table 4:** Subgroup optimized parameter values compared to Smith *et al*. model based norms.

| Subgroup | $E_{lv}$ | $\lambda_{lv}$ | $E_{rv}$ | $\lambda_{rv}$ | $E_{pa}$ | $E_{pu}$ | $R_{pul}$ | $E_{sa}$ | $R_{sys}$ |
| --- | --- | --- | --- | --- | --- | --- | --- | --- | --- |
| HFrEF | 0.5 | 1.1 | 2.5 | 1.1 | 5.2 | 7.3 | 0.7 | 1.5 | 1.5 |
| HFpEF 1 | 0.6 | 1.0 | 1.7 | 1.0 | 3.1 | 9.4 | 0.6 | 1.3 | 0.8 |
| HFpEF 2 | 2.1 | 2.2 | 2.2 | 1.7 | 3.2 | 7.5 | 2.0 | 2.6 | 1.7 |
| NCC | 1.4 | 1.7 | 2.8 | 0.9 | 1.9 | 5.3 | 1.9 | 1.4 | 1.1 |

HFrEF – heart failure with reduced ejection fraction. HFpEF – heart failure with preserved ejection fraction. NCC – not consistently clustered.

**Figure 7.**
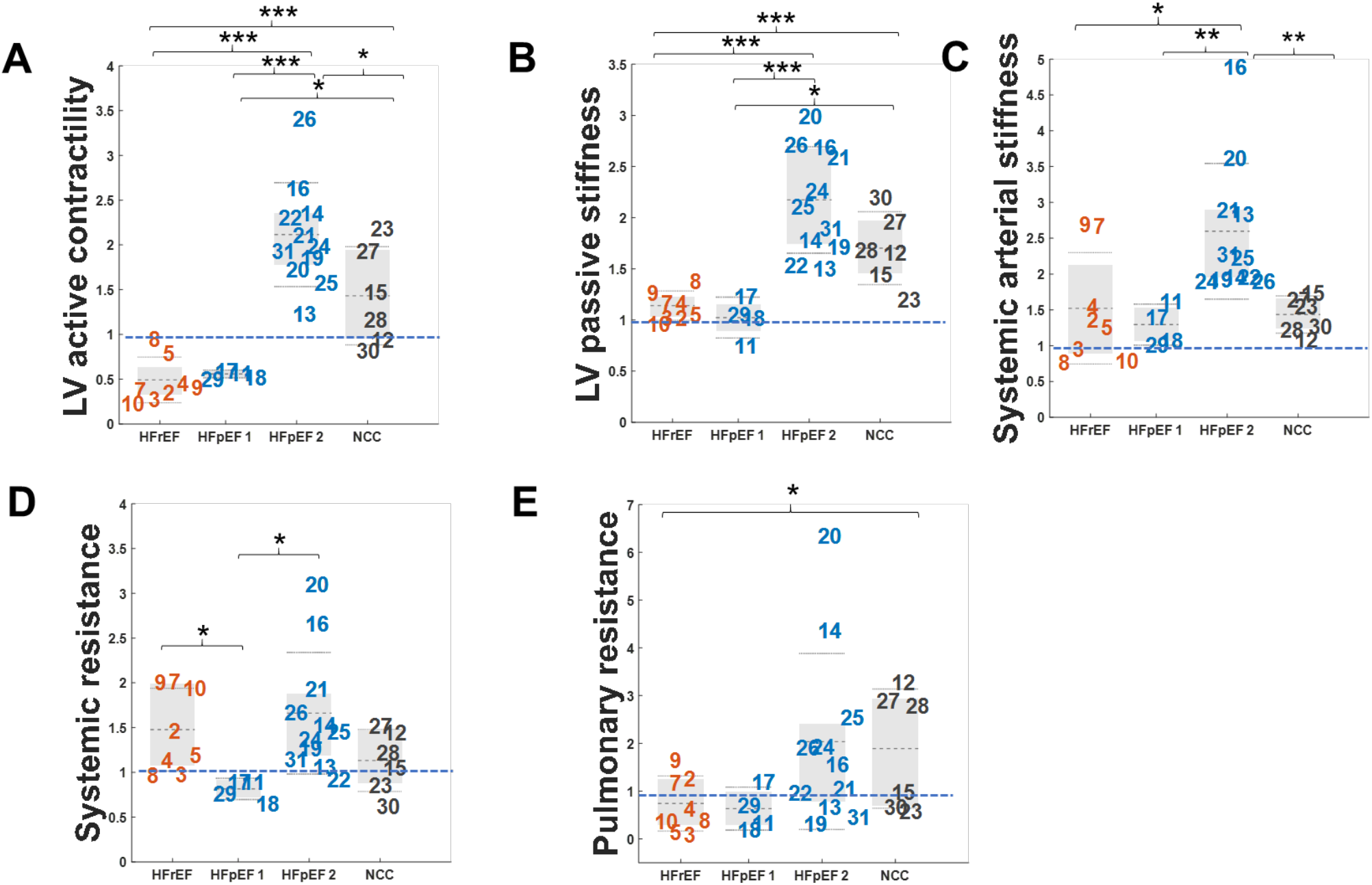
Analysis of our optimized parameters gives us an understanding of the mechanistic differences between the three HFpEF groups which cannot be seen by analyzing the clinical measures alone. **A-E** Left ventricular active contractility, left ventricular passive stiffness, systemic arterial stiffness, systemic and pulmonary resistance respectively relative to the normal Smith model parameter values indicated in blue dashed line. Gray dashed line is average, light dotted is standard deviation and the grey box contains the middle 50% of the parameter values (* p-value <.05, ** p-value] <.01, *** p-value <.001).

Similar to LV active contractility, when comparing LV passive stiffness **(Figure 7B)** no significant differences were observed between HFrEF and HFpEF Group 1. Both groups had near normal values when compared to model-based norms unlike HFpEF Group 2 and NCC which were found to be above threshold **(Table 4)**. When compared to both HFrEF and HFpEF Group 1, the HFpEF Group 2 shows significantly higher LV passive stiffness (p-value < 0.001). Likewise, the LV passive stiffness in the NCC subgroup is significantly higher (p-value < 0.001 and p-value < 0.05, respectively) when compared to HFrEF and HFpEF Group 1. No significant differences were found between HFpEF Group 2 and NCC in this parameter.

Looking at the parameter representing systemic arterial stiffness **(Figure 7C)**, all groups are found to be above threshold when compared to model-based norms **(Table 4)**. When compared to both HFrEF and HFpEF Group 1, the HFpEF Group 2 shows significantly higher systemic arterial stiffness (p-value < 0.05 and p-value < 0.01, respectively). Just like when comparing LV active contractility and passive stiffness no significant differences were observed between HFrEF and HFpEF Group 1 in this parameter, yet despite having similar trends HFpEF Group 2 had a significantly higher (p-value < 0.01) systemic arterial stiffness when compared to the NCC group.

Strikingly, systemic resistance **(Figure 7D)**, in all groups except HFpEF Group 1 is found to be above threshold when compared to model-based norms **(Table 4)**. This is the only parameter in which significant differences are observed between HFrEF and HFpEF Group 1 (p-value < 0.05). When compared to HFpEF Group 1, HFpEF Group 2 shows significantly higher systemic resistance (p-value < 0.05). No significant differences appear between HFrEF, HFpEF Group 2 and NCC. Of note, significantly higher (p-value < 0.05) pulmonary resistance **(Figure 7E)** was found in the NCC group, when compared to the HFrEF group. Pulmonary resistance was almost reduced to half compared to the model-based norm in HFrEF and HFpEF 1 groups, while it was almost doubled in the HFpEF group 2 and NCC **(Table 4)**.

Our results show that the main mechanistic parameter driving both HFrEF and HFpEF Group 1 is reduced left ventricular contractility as well as left ventricular passive stiffness **(Figure 7A, B)**, indicating that systolic dysfunction is the primary driver for both patient cohorts. Consistent with the classical definition of HFpEF being characterized by diastolic dysfunction, our simulations show that our HFpEF Group 2 has increased left ventricular passive stiffness, as well as increased left ventricular active contractility at rest **(Figure 7A, B)**. Moreover, HFrEF and all HFpEF groups show systemic arterial stiffness relative to model-based norms **(Figure 7C, Table 4)**. Likewise, HFrEF and all HFpEF groups (except HFpEF Group 1) have higher systemic resistance relative to model-based norms **(Figure 7D)**. Taken together, these results stress that the systemic vasculature plays a role in heart failure regardless of HF type.

Using the four groups determined from the model parameter optimization and clustering techniques, we again compared the patient RHC and TTE clinical data measurements **(Figure 8)**. Ejection fraction between the HF groups reveals very significant differences between all our distinct HF subgroups **(Figure 8A)**. As expected, the ejection fraction in HFpEF Group 1 is significantly higher (p-value < 0.05) than that of our HFrEF cohort. Likewise, the ejection fraction in HFpEF Group 2 and NCC is very significantly higher (p-value < 0.001) than the HFrEF cohort. Of note, the different HFpEF groups have an ejection fraction above 50%, consistent with their HFpEF diagnosis yet significant differences amongst ejection fraction between HFpEF subgroups are observed. Although no significant differences were found between HFpEF Group 2 and the NCC group, these two groups have significantly higher ejection fraction (p-value < 0.01 and p-value < 0.05, respectively) than HFpEF Group 1.

**Figure 8.**
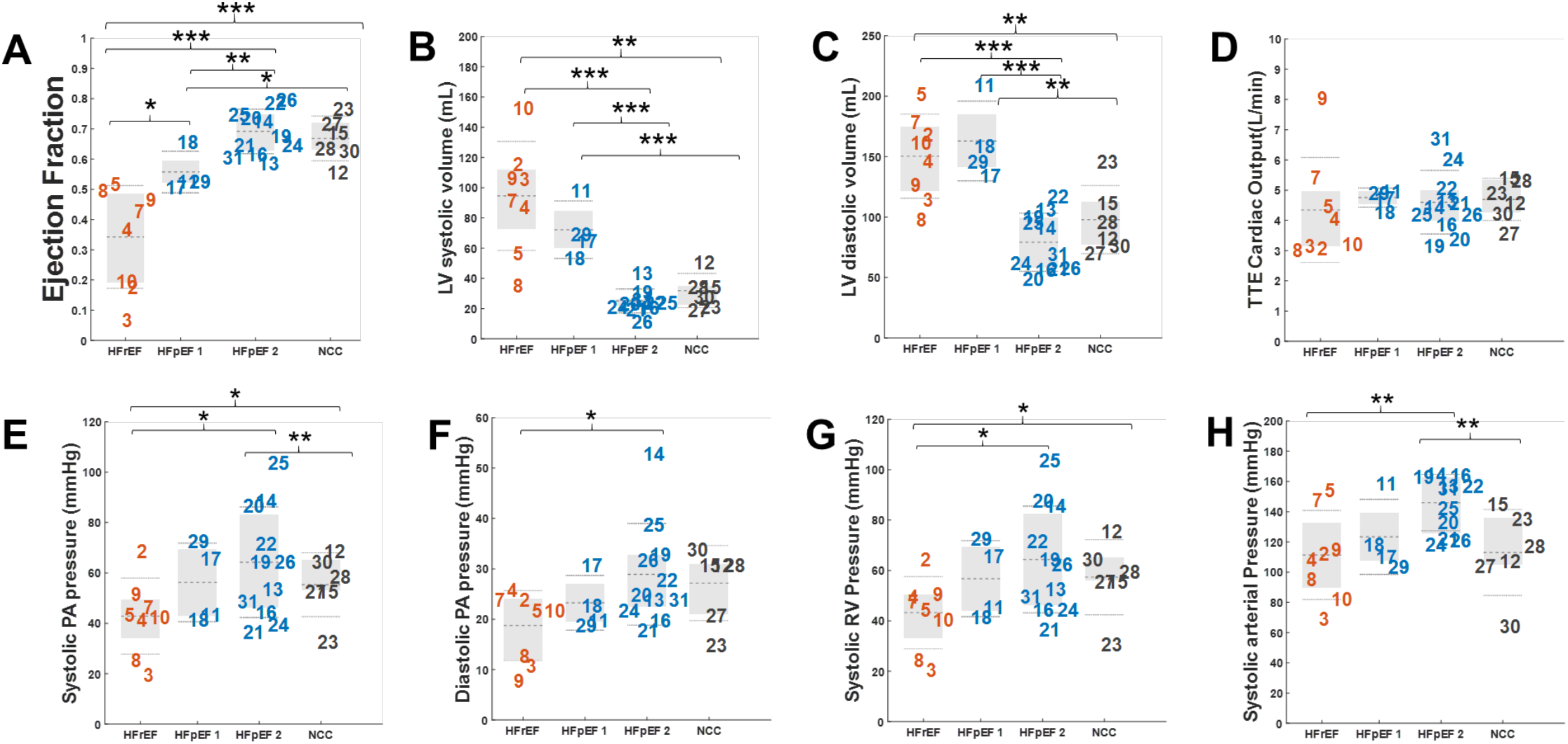
Significant differences are seen in TTE and RHC clinical data between Heart Failure patients based on novel classification based on clustering analysis of model-based optimized parameters. Gray dashed line is average, light dotted is standard deviation and the grey box contains the middle 50% of the clinical values (* p-value <.05, ** p-value <.01, *** p-value <.001).

HFrEFs shows significant higher left ventricular systolic volumes **(Figure 8B)** when compared with the HFpEF Group 2 and NCC groups (p-value < 0.01 and p-value < 0.001, respectively), yet no significant difference was found between HFrEF and HFpEF Group 1. Similar to the HFrEF group when comparing HFpEF Group 1 to HFpEF Group 2 and NCC groups, it showed significant (p-value < 0.001) higher left ventricular systolic volumes than both groups. No significant differences in left ventricular systolic volume were found between HFrEF and HFpEF Group 1. Left ventricular diastolic volumes showed the same trend as volumes in systole, no significant differences were found between HFrEF and HFpEF Group 1 **(Figure 8C)**. Likewise, HFrEFs showed significant higher diastolic volumes than HFpEF Group 2 and NCC (p-value < 0.001 and p-value < 0.01, respectively). HFpEF Group 1 had significantly higher diastolic volumes than HFpEF Group 2 and NCC (p-value < 0.001 and p-value < 0.01, respectively). No left ventricular diastolic volume differences were found between HFpEF Group 2 and NCC. Despite this, TTE cardiac output did not reveal any differences between groups **(Figure 8D)**. On the other hand, CO in the RHC measurements did reveal significant differences (p-value < 0.05) between the HFpEF Group 1 and HFrEF groups (**Supplemental Figure 8G**).

RHC pressure measurements revealed significantly higher pulmonary arterial systolic pressures in the HFpEF Group 2 when compared to HFrEF and NCC group (p-value < 0.05 and p-value < 0.01, respectively). The NCC group also showed significant higher (p-value < 0.05) pulmonary arterial systolic pressure when compared to HFrEF (**Figure 8 E**). Similar to its behavior during systole, HFpEF Group 2 showed a significant (p-value < 0.05) higher pulmonary arterial diastolic pressure when compared to the HFrEF group (**Figure 8 F**). RHC pressure measurements likewise revealed higher right ventricular systolic pressures (p-value < 0.05) in HFpEF Group 2 when compared to HFpEF Group 1 as well as when comparing the HFpEF NCC group to HFrEF (**Figure 8 G**). Systolic arterial pressure in HFpEF Group 2 was also found to be significantly higher (p-value < 0.05) when compared to HFrEF and NCC groups.

These results show that analysis of data from RHC and TTE of heart failure patients using a closed loop model of the cardiovascular system is able to identify key parameters representing hemodynamic cardiovascular function in HFrEF and HFpEF patients. Analyzing these optimized parameters representing cardiovascular function with the aid of machine learning techniques gives us an understanding of the mechanistic differences between three different HFpEF groups that is not seen analyzing clinical measures alone.

## Discussion

Our analysis of optimized parameters representing patient specific cardiovascular mechanics coupled with unsupervised machine learning techniques enabled us to identify distinct HFpEF groups that share similar deep mechanistic phenotypes. These HFpEF patient groups cannot be discerned from the clinical data alone. Moreover, the mechanistic parameters used to distinguish HFpEF groups, allowed us to describe not only the functional details of the cardiovascular system for each population but for each patient in the population. The approach to represent cardiovascular mechanics in this study not only considers hemodynamics in the heart, but also the pulmonary and systemic vasculature providing a more holistic view of the cardiovascular state for each population and each patient.

### Clustering of HFpEF groups

While HFrEF is characterized by a well-defined phenotype, HFpEF comprises a large variability in the constellation of changes at the cardiovascular system level. We found that the HFpEF group presented here can be subdivided into 3 subcategories: HFpEF group 1 described as “HFrEF-like HFpEF”, HEpEF group 2 as “consistent HFpEF”, and a final NCC as “HFpEF group that was not consistently clustered” (**Figure 6F)**. Using PCA and clustering techniques to analyze clinical data alone, the same HFpEF distinctions cannot be seen (**Figure 4D**) suggesting that key discriminators of HFpEF into distinct phenotypes reside at the mechanistic level revealed only by using our patient-specific computational model. Simply looking at the underlying mechanistic parameters from our patient-specific modeling (**Figure 7** for groups HFpEF 1, 2 and NCC), we see that the range of values for the HFpEF population was widely heterogeneous. After finding the 2-dimensional reduced space of parameters derived from the patient-specific tuned models that produces the largest variation across patients through PCA, we can see that there are some HFpEF patients that lie in the same region as the HFrEF patients (**Figure 7A)**. Extracted physiological parameters, such as left ventricular active contractility and left ventricular passive stiffness, were shown to play an important role in describing these distinct patient populations.

### HFpEF Group 1 as HFrEF-like HFpEF

In our HFrEF population, we see a consistent normal left ventricular passive stiffness, coupled with a reduced maximal contractility at rest in the left ventricle (**Figure 7A**,**B**). This is consistent with the current understanding of HFrEF, where systolic dysfunction is the main pathological characteristic describing this phenotype(Pinilla-Vera *et al*., 2019). In our heterogenous HFpEF population, we surprisingly found that HFpEF group 1 (HFrEF-like HFpEF) shares the same mechanistic parameter trends as the HFrEF group except lower systemic resistance (p<0.05 and near normal, **Table 4**). Of note when looking at the clinical data comparison between our new HF groups, patients in the HFpEF group 1 have significantly higher ejection fraction than HFrEFs but a significantly lower ejection fraction than the other two HFpEF groups (**Figure 8A**). This could be explained by the fact that while both HFrEF and HFpEF group 1 show systolic and diastolic left ventricular volume overload when compared to HFpEF2 and NCC groups, the HFpEF group 1’s systolic volume is lower than that of the HFrEF group (**Figure 8 B, C**). These results shed some hope at the possible treatment options for these HFpEF patients. Since they share such similar physiological characteristics with the HFrEF cohort, therapeutic strategies currently employed to alleviate systolic dysfunction in HFrEF patients might be employed in other patients similar to HFpEF Group 1. Likewise, our results indicate that both HFrEFs and HFpEF group 1 patients would benefit from treatments that would improve left ventricular contractility.

### HFpEF Group 2 and NCC

The second or “pure” HFpEF group has very high left ventricular passive stiffness, coupled with an increase in maximal contractility at rest in the left ventricle (**Figure 7A, B, Table 4**). Although these patients maintain a higher ejection fraction than the other groups their reduced ability to fill during diastole leads to very low systolic volumes (**Figure 8 B, C**). This would be particularly challenging in situations where recruiting a higher stroke volume is necessary, such as during exercise. The elevated ventricular stiffness in this cohort could explain the increased levels of systolic and diastolic pulmonary arterial pressure, systolic right ventricular pressure and arterial pressure observed in the clinical data of these patients (**Figure 8E-H**). The combination of increased left ventricular passive stiffness, systemic arterial stiffness and higher pressures in the pulmonary and systemic vasculature may account for the increased pulmonary and systemic resistance also observed in this patient cohort (**Figure 7C-E**).

Our NCC group, created out of need to cluster patients that were distinct from HFrEFs but did not fall clearly into the HFpEF group 1 or 2 categories, as it falls between groups it represents a “spectrum” of patients more than a clearly defined subgroup, perhaps this is a population of HFpEF undergoing remodeling and through time decompensating into HFrEF. It clearly does not behave like a HFrEF or HFpEF group 1 as it shows high left ventricular passive stiffness, coupled with an increase in maximal contractility at rest in the left ventricle (**Figure 7A, B, Table 4**). Yet whatever stiffness or increased in left ventricular contractility it has; it still shows a milder phenotype than that of the “pure” HFpEF group 2. Looking at the clinical data, NCC’s ejection fraction as well as systolic and diastolic left ventricular volumes are like the ones found in HFpEF group 1 patients (**Figure 8 A-C**). The differences in physiological mechanistic behavior is perhaps accounted to the lower systemic and pulmonary pressures observed in the NCC group when compared to HFpEF Group 2 (**Figure 8E-H**).

### Possible Clinical presentation of HFpEF subgroups

The distinct HFpEF populations found here are consistent with the recent studies describing HFpEF as a disparate phenotype. In one such study, analysis of RNA sequencing of right ventricular septal endocardial biopsies on control, HFrEF, and HFpEF patients through unsupervised machine learning identified three HFpEF transcriptome subgroups with distinctive pathways and clinical correlations(Hahn *et al*., 2021). These HFpEF subgroups included: a hemodynamic-driven HFpEF group close to HFrEF showing the worst clinical outcomes; a HFpEF cohort with smaller hearts and inflammatory and matrix signatures; and a third heterogeneous phenotype with worse HF symptoms but lower NT-proBNP and smaller hearts that remains distinct from HFrEFs. Patients in the HFpEF Group 1 of the Hahn *et al*. study also had higher LV dimensions, perhaps consistent with the ventricular volume overload we observed in both HFrEF and HFpEF group 1 patients in our study. Since the transcriptome of this Group 1 in which mitochondrial adenosine triphosphate synthesis and oxidative phosphorylation are downregulated is also closest to HFrEF, the authors suggests therapies targeting the latter molecular mechanisms may benefit this HFpEF subgroup as well. Of note HFpEF Group 2 in the Hahn *et al*. study was all female, had the smallest LV size, lowest NT-proBNP, but higher number of CD68+ inflammatory cells. Perhaps the small LV size seen in these patients would be in accordance to the very small LV volumes observed in our HFpEF group 2 patients, the only group in our cohort in which the majority of the patients were also female. Likewise, our NCC group could belong to the third heterogeneous phenotype with worse HF symptoms but lower NT-proBNP and smaller hearts that remains distinct from HFrEF. Based on their transcriptome results, the authors suggest that therapies targeting protein processing, metabolism, inflammation, or matrix remodeling may more benefit their HFpEF groups 2 or 3; while obesity reduction would likely help all groups (all patients groups in our study had an average BMI>30, indicating that they likewise would benefit from obesity reduction).

Another study utilizing quantitative echocardiography phenotyping with unsupervised machine learning identified 3 HFpEF phenogroups with differing clinical and echocardiographic characteristics and outcomes: one group with natriuretic peptide deficiency syndrome; a second group with extreme cardiometabolic syndrome; and a third group with right ventricular cardio-abdomino-renal syndrome (Shah, 2019). Amongst one of the characteristics of the second HFpEF phenogroup in this study was that it had the most severely impaired cardiac relaxation compared to the other HFpEF groups. Our second or “pure” HFpEF group showing very high left ventricular passive stiffness perhaps falls in this same category.

The studies by both Hanh *et al*. and Shah *et al*. show novel classifications of HFpEF subgroups based on transcriptomic analysis of endomyocardial biopsy obtained through right heart catheterization and a detailed clinical, laboratory, ECG, and echocardiographic data phenotyping respectively. While descriptive and pointing out clinical markers that may describe these novel HFpEF classifications (i.e. NT-proBNP marker, inflammatory signal differences between groups) the nature of cardiovascular hemodynamics and its relationship with the pulmonary and systemic vasculature defining HFpEF subgroups and the uniqueness of each patient within a group requires a deep phenotyping approach using clinical data to power cardiovascular model-informed machine learning.

### Role of the systemic vasculature in heart failure

The physiological parameters derived from our cardiovascular system mode aligns with the understanding that HFrEF patients have a normal LV passive stiffness, reduced LV contractility, increased systemic resistance, and slightly increased arterial stiffness as well as near normal pulmonary resistance when compared to normal cardiovascular function (**Figure 7**). This alignment of the underlying mechanistic cardiovascular parameters of the model with the conventional wisdom concerning HFrEF suggests that the clinical data used here is sufficient to describe HFrEF. This also gives us confidence in the profile of the deep phenotypes of HFpEF that are revealed here. This also suggests that focusing purely on cardiac function may consistently capture the underlying dysfunction in HFrEF but is not a good approach to understanding HFpEF. For example, HFpEF Group 2 patient 20 has some LV hypercontractility and increased LV passive stiffness but exhibits large differences in the systemic and pulmonary vasculature. Likewise, NCC patient 12, shows increased LV passive stiffness and increased pulmonary resistance but without displaying the LV hypercontractility or high levels of systemic resistance and arterial stiffness other HFpEF group 1 or NCC patients show. In both patients, addressing the cause for increased resistances in the systemic and pulmonary vasculature may aid reduce the burden the heart has set up to compensate when HF is present

### Limitations

In this study, a general HFrEF group was used as the only reference HF type. For a larger patient cohort applying a thorough clustering analysis with not just HFpEF and HFrEF phenotypes but also other HF diagnoses might shed even greater clarity into the physiological differences between groups. Based on the TTE systolic and diastolic volumes HFrEF patients 1 and 6 exhibit volume overload and seem to suffer from severe ventricular dilation. Our cardiovascular systems model was unable to account for these large volumes. Hence, making appropriate changes recruiting more stressed volume in the model, decreasing ventricular elastance, or implementing a more detailed model may capture the pathological complexity of these patients. Regardless of future directions taken, the physiological parameters derived from the simple cardiovascular system model can still be useful determinants for HF classification purposes beyond ejection fraction.

## Conclusions

HFrEF and HFpEF have classically been defined based on ejection fraction. The HFrEF diagnosis itself is much more understood than HFpEF, which is largely heterogeneous. In accordance with other recent studies, we have determined 3 subgroups of HFpEF with our methodological deep phenotyping approach that uses cardiovascular model-informed machine learning: a HFrEF-like HFpEF group, a “pure” HFpEF group, and a group that exhibits characteristics of both. Moreover, these subgroups could not be distinguished based on the clinical data alone. Ultimately, the combination of mathematical modeling analysis and machine learning techniques provides immense insight into the classifications of heart failure as a pathology.

